# Quantifying Expansion Microscopy with DNA Origami Expansion Nanorulers

**DOI:** 10.1101/265405

**Authors:** Max B. Scheible, Philip Tinnefeld

## Abstract

In the past decade super-resolution microscopy^1,2^ developed rapidly and allowed seeing new structural details in fluorescence microscopy, especially in the field of bioimaging^3-6^. Most of the evolving techniques like (d)STORM^7,8^, STED^9^ or SIM^10^ – just to name a few – focused on overcoming the diffraction limit^11^ by increasing the technical resolution of the microscope, often in combination with specifically designed probes, for example by using blinking fluorescent dyes^7,8,12,13^. But the approach to improve the effective microscope resolution of a system is not the only way to reveal new insights. An alternative approach is based on physically expanding the sample to increase its size by a multiple and to subsequently image the sample by conventional fluorescence microcopy.^14^ The technique termed expansion microscopy (ExM) allows effective resolution below the diffraction limit and therefore directly complements and competes with established super-resolution techniques, but offers the advantage to use standard confocal or wide-field microscopes.

The idea behind ExM is to embed the sample – for instance tissue or fixed cells – in an electrolytic polymer and to expand the gel by dialysis to gain a physical magnification. To visualize the target of interest, the target is labeled with an antibody-fluorophore complex, which is further cross-linked to the polymer before expansion^14,15^. After digesting the sample, the fluorophore is still linked to the polymer mesh, mimicking shape, structure and dimension of the original target structure. Subsequent expansion of the polymer then allows diffraction-limited imaging with converted resolutions down to 70 nm. In the original work, labeling was carried out by a primary antibody and a DNA labeled secondary antibody. The DNA strand served as adapter for a complementary DNA strand that carries both functionalities, i.e. the fluorescent dye and the reactive acrydite group for connecting to the polymer. Variations of the original expansion microscopy were demonstrated using for instance RNA label in combination with FISH^16^ or completely avoiding nucleotides by direct-labeling via proteins^15^. Usually these techniques reach macroscopic expansion factors (EF_ma_) of 3-5 (the factor of size increment of the side length of the gel before and after expansion).^14-16^ The expansion factor can be improved by applying different electrolytic gels (EF_ma_ up to 10)^17^ or by using an approach called iterative expansion microscopy^18^. Here, two successive gels are applied to the same sample reaching an EF_ma_ of up to 20, which allows resolutions down to 25 nm. Alternatively higher resolutions can be achieved by combining ExM with other super-resolution techniques like structured illumination microscopy (SIM) to image cellular components with a resolution down to 30 nm.^19^

Generally, the magnification achieved by expanding the gel is quantified by consulting the macroscopic swelling of the gel. It currently remains an open question and a challenge to objectively evaluate whether the macroscopic swelling of the gel homogeneously correlates with the expansion of target structures on the microscale. Is the expansion isotropic and homogeneous? Do surface effects and breaks play a role? Ultimately, the resolution improvement of expansion microscopy should be limited by the mesh size of the polymer matrix. Could this limit be visualized?

To quantify the microscopic expansion factor (EF_mi_) – instead of the macroscopic one – a biocompatible soft matter with defined properties is required, so that the expansion can be tracked and evaluated from the inside of the gel. Here the advantages of DNA nanotechnology come into play. In the past decade, nanoscale structures made from DNA conquered the field of biotechnology^20^, especially due to the development of the DNA origami technique^21^. This technique allows building of arbitrary two- and threedimensional shapes with defined dimensions of up to a few hundred nanometers^22,23^, a size range, which was not accessible by biosynthetic approaches before. The ability to precisely modify these structures at will by linking molecules to the DNA at defined positions enabled a completely new type of functional nanomaterial.^24-27^ Using fluorophores as specific markers on the nanostructure and separating these fluorescent marks with a defined distance creates a new class of measurement standards: the DNA nanorulers.^28-30^

In this work we transfer the concept of DNA origami nanorulers to expansion microscopy to quantify the magnification of the sample in 2D and to determine the microscopic expansion factor EF_mi_. To this end, we modify a nanoruler with 160 nm spacing between two marks in a way that we can crosslink the dye labels to the polymer network and expand it. We demonstrate that a structure with a defined pattern below the diffraction limit can be imaged with a conventional fluorescence microscope by means of ExM. DNA nanorulers will enable to resolve the open quantification questions of expansion microscopy and could even be applied as in situ measuring reference samples.

The expansion nanoruler is a modified version of a GATTA-SIM 160R nanoruler labeled with ATTO 647N dyes (Fig. 1a). On this DNA origami platform two areas are specified, so-called marks, with a center-to-center distance of 160 nm and a mark size of ~20 nm (Supplemental Fig. S1 and Fig. S2). Each mark consists of twenty single-stranded staple extensions, wherein each extension is a 21 nucleotide long single DNA strand with a designed sequence S1’. S1’ extensions can be labeled with complementary S1 strands, which fulfill a twofold purpose. First, an ATTO 647N dye is conjugated to the 3’-prime end of the S1 strand, which enables imaging under pre- and post-expansion conditions. Second, the 5’-prime end is labeled with an acrydite group to covalently bind to the polymer matrix during gelation.

**Figure 1:**
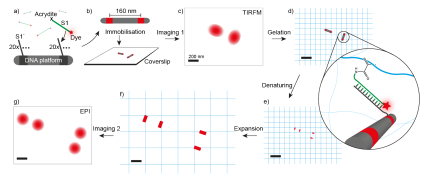
Procedure for expanding DNA nanorulers in 2D. a) S1 strands are labeled with fluorescent ATTO 647N dyes and acrydite groups. They can hybridze to complementary S1’ strands protruding from a DNA origami platform (nanoruler). The platform provides two separated marks with twenty S1’ extensions, respectively. b) The nanorulers carrying two marks separated by 160 nm are immobilised on a BSA/neutravidin surface. c) Diffraction-limited imaging can not resolve the two marks and shows them as a single overplapping PSF. d) By adding an electrolytic gel the acrydite of the S1 strand crosslinks to the acrylamide during the polymerization process. e) Denaturing of the DNA separates the S1 strand from the nanoruler while the strand maintains its position linked to the polymer matrix and subsequently mimics the marks of the ruler. f) Expansion of the gel increases the distance between the two marks. g) Epi wide field imaging displays the two marks of the expansion nanoruler without overlapping PSFs.

The expansion nanoruler is additionally equipped with biotin anchors on the bottom side of the platform to immobilize it on a BSA/neutravidin surface (Fig. 1b). Single nanorulers are then imaged on a single-molecule sensitive wide field or TIRF setup (Fig. 1c). Since the nanoruler displays a distance of 160 nm (Supplemental Fig. S2), which is well below the diffraction limit of ~283 nm (Supplemental Information), the point spread functions (PSFs) of each mark overlap and every nanoruler appears as one single spot. Embedding the sample in an actively polymerizing electrolytic polyacrylamide gel leads to covalent binding of the nanoruler to the surrounding polymer network via the acrydite modification, so that the S1 strands are connected between the immobilized nanorulers and the established gel matrix (Fig. 1d).

By a denaturing step the DNA origami platform is decomposed and subsequently the DNA duplex between nanoruler and the S1 strands is dissociated (Fig. 1e). The fluorescently labeled S1 strands mimic the positions of the two marks on the nanoruler and are simultaneously bound to the gel. Linear expansion of the gel increases the distance between the two marks leaving the diffraction-limited regime behind (Fig. 1f). Subsequent imaging displays the two separated marks even with conventional microscopes (Fig. 1g).

Experimentally, the gel was expanded on top of a microscope coverslip. To improve the plain attachment to the coverslip the gel was loosely covered with a second coverslip. The ratio of the size of the gel before and after expansion (Fig. 2a and b) yields the macroscopic expansion factor *EF_ma_.* Taking the values from Figure 2a and b the expansion factor is EF_ma_ = 3.6. Using the specifications of the expansion nanoruler with an intermark distance of *d_pre_* = (160 ± 9) nm (Supplemental Fig. S2) we would expect a post-expanded distance d’_*post*_r = 3.6 * d_*pre*_ = (576 ± 32) nm.

**Figure 2:**
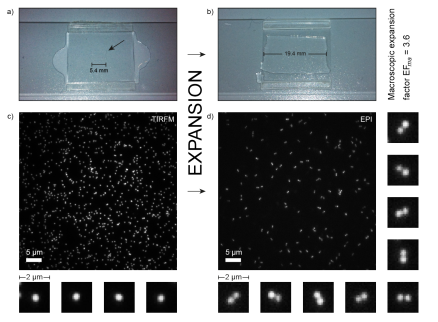
The electrolytic polyacrylamide gel before expansion in between two coverslips with an average width of 5.4 mm. b) The same gel after expansion with an average width of 19.4 mm. The macroscopic expansion factor is 3.6. c) TIRFM image of immobilized nanorulers before gelation and expansion carrying ATTO 647N dyes. Nanorulers with 160 nm intermark distances appear as one spot due to the diffraction-limited imaging (selected zoom-ins). d) After expansion nanorulers are imaged in epi-fluorescence and are clearly resolved, represented by two adjacent spots (selected zoom-ins).

Imaging of the pre-expanded nanorulers in TIRF mode shows a homogeneous distribution of single spots, which originates from the overlapping PSFs of the two marks on the expansion nanoruler (Fig. 2c). The surface density of immobilized structures is adequate, showing many single structures with a narrow intensity distribution (Supplemental Fig. S3) and rather few overlaying structures/clusters. Selected single structures show spherical PSFs with an average FWHM of (363 ± 8) nm. The substructure with 160 nm spacing is therefore too small to shape the superposed PSFs into an elliptic contour.

After expansion nanorulers are visible in the gel roughly 1 μm above the coverslip surface and are therefore imaged in EPI mode (Fig. 2d). It is clearly observable that there is a vanishing number of single spots but pairs of two adjacent spots displaying the expanded nanoruler. The density of structures is reduced in accordance with the volume increase of the sample. This change in structural density can provide a first hint towards a quantitative analysis of the microscopic expansion. Before expansion five images with n_*pre*_ = 3398 structures in total are analyzed and yield a structural density of (2.59 ± 0.15) structures per 10 μm^2^. After expansion this density drops to (0.37 ± 0.04)/10 μm^2^ (eleven images with n_*post*_ = 1077 structures) resulting in a rough estimate for the microscopic expansion factor with EF_*mi*_(density) ≈ 2.6. The clear presence of two-spot structures in combination with the drop in structural density shows that DNA origami nanorulers initially bound to a coverslip can be quantitatively and homogenously expanded and are therefore applicable as nanoscopic expansion rulers.

For further quantification, the expansion nanorulers are analyzed using the GATTAnalysis software. GATTAnalysis uses an automated spot finder and fits two 2D-Gaussians to each structure to determine the center-to-center distance and the FWHM of the spots. The distance analysis of *n_post_* = 1077 structures yields an average distance of d*_post_* = (445 ± 56) nm (Fig. 3a). Thus the microscopic expansion factor referring to the total number of expanded nanorulers can be calculated to EF_mi_(total) = *d_post_/d_pre_* = 2.8. The size of the PSF of each spot is (351 ± 18) nm and therefore similar to the FWHM values in the pre-expanded state (Fig. 3b).

**Figure 3:**
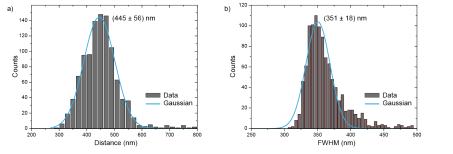
A) Quantitative analysis of n = 1077 nanorulers yields an average ruler distance of (445 ± 56) nm after expansion. b) The FWHM after expansion is (351 ± 18) nm. Results were obtained by an automated analysis using the GATTAnalysis software.

To investigate the homogeneity of the expansion the eleven single images were individually analyzed resulting in individual expansion factors (Supplemental Information). The microscopic expansion factor was homogeneous over a large part of the coverslip and different regions yield a factor of EF_*mi*_(individual) = 3.0 ± 0.1. Furthermore, we could not detect any breaks or cracks of the gel but it appeared homogeneous in the full area investigated.

It turns out that the microscopic expansion factor deviates from the macroscopic one and it is slightly smaller. Apart from that different approaches to determine the microscopic expansion factor show additional deviations between the approaches. The origin of the deviation of macroscopic and microscopic expansion factor is not straightforward. In first approximation, the microscopic expansion factor is based on the properties of the DNA nanorulers. The validity to use nanorulers as benchmark structures and traceable measurement standards has recently been shown by Raab et al. who verified their usefulness as reliable and quantitative tool.^31^ The covalent binding to the polymer matrix with subsequent expansion, however, introduces various imponderables, which can directly influence the measured intermark distances. We see two main possibilities for the different microscopic and macroscopic expansion factors. First, the expansion rulers are initially placed at the coverslip surface. The swelling occurring directly at the interface between gel and coverslip might be reduced by friction yielding less expansion close to the coverslip. Second, the nanorulers might be tilted during expansion which is not detected in our 2D imaging and analysis. At this point, we cannot decide what the true origin of the deviation is. Both of the discussed error sources considered represent a sort of deviation from ideal isotropic expansion behavior, which deserves further attention and indicates the need to quantitatively and microscopically determine expansion factors.

The fact that the microscopic expansion factor shows slight deviations itself – with regards to the analysis method, which is used – is helpful when it comes to the interpretation of the data. The results for EF_mi_(density) obtained from the structural density analysis are only a rough estimate but already get close to the other expansion values and therefore illustrate an approach, which is fast and qualitative to retrieve. Interestingly, this value even shows the smallest expansion factor. This is a very good hint that the binding efficiency of nanorulers to the gel matrix is very high, since unbound structures would be digested and not expanded and subsequently lead to a lower structural density after expansion. Low structural densities would induce a bias towards larger expansion factor values, which cannot be found in this case. Consequently, a small expansion factor EF_mi_(density) strengthens the claim of a very high coupling yield.

The difference between the microscopic expansion factor EF_mi_(total) (using the total number of nanorulers) compared to the individual approach with EF_mi_(individual) (to explore the homogeneity of the expansion) can be traced back to different statistical approaches to evaluate the data (Supplemental Information). However, EF_mi_(individual) yields a very small standard deviation of only 0.1 proving the homogeneous expansion over a large field of the coverslip.

In this work, we showed that DNA origami nanorulers that had proven their potential for several super-resolution techniques already^29,30,32^ can be especially valuable tools for the microscopic characterization of magnification and resolution in expansion microscopy. We found a substantial deviation of microscopic and macroscopic expansion factor which should be considered for the quantitative discussion of expansion microscopy data of biological cells and tissues. Beside that we showed that different approaches to determine the microscopic expansion factor can deviate from each other – a fact which should be taken into account and discussed in detail when interpreting the quantitative expansion of the target. The introduced expansion nanorulers present a powerful tool to reveal the true origin of anisotropy and heterogeneity in expanded microscopy specimen. As the expansion nanorulers undergo the same coupling and expansion process as the biological target structure the found deviations of expansion factors should be representative for the biological target as well.

The shown approach helps to develop the expansion microscopy technique from an imaging application towards a more quantitative evaluation of imaging data. 3D studies will be a key to implement the nanoruler tools directly into cell/tissue imaging and to combine the biological question with quantitative conclusions. Since the proof-of-principle is demonstrated a new field opens up with the potential to answer questions with respect to quantitation of microscopic magnification, the anisotropy or local dependence of magnification, or the binding efficiency of dyes to the gel. Ultimately, the maximum possible resolution of expansion microscopy related to the density of labeling the mesh size in the polymer might be unveiled. These conclusions call for *in situ* expansion nanorulers for taking expansion microscopy to a quantitative level.

## Acknowledgements

M.B.S. and P.T. thank E. Boyden (Massachusetts Institute of Technology), K. Weisshart (Carl Zeiss Microscopy GmbH), J.J. Schmied and C. Forthmann (both GATTAquant GmbH) for helpful discussions and T. Dammeyer (TU Braunschweig) for providing the scaffold DNA. Further, we acknowledge the support of the Technische Universitaet Braunschweig, the Institut fuer Theoretische und Physikalische Chemie, the Braunschweig Integrated Center for Systems Biology (BRICS), and the Laboratory for Emerging Nanometrology (LENA). The work was also supported by GATTAquant GmbH. P.T. gratefully acknowledges support by the DFG (INST 188/401-1 FUGG and excellence clusters Nanosystems Initiative Munich (NIM) and CIPSM (Center for Integrated Protein Science Munich)). This project was also supported by the European Union’s Horizon 2020 research and innovation program under grant agreement No 737089 (Chipscope).

## Author contributions

M.B.S. planned, conducted and evaluated the experiments; M.B.S. and P.T. wrote the manuscript.

## Additional Information

Supplementary Information is available.

